# Coordination of multiple joints increases bilateral connectivity with ipsilateral sensorimotor cortices

**DOI:** 10.1101/656819

**Authors:** Kevin B. Wilkins, Jun Yao

**Affiliations:** Department of Physical Therapy and Human Movement Sciences, Northwestern University, 645 N Michigan Ave, Suite 1100, Chicago, IL 60611, USA; Northwestern University Interdepartmental Neuroscience, Northwestern University, 320 E. Superior St, Chicago, IL 60611, USA; Department of Biomedical Engineering, Northwestern University, 2145 Sheridan Road, Evanston, IL 60208, USA

**Keywords:** electroencephalography (EEG), dynamic causal modeling (DCM), connectivity, motor planning, complexity

## Abstract

Although most activities of daily life require simultaneous coordination of both proximal and distal joints, motor preparation during such movements has not been well studied. Previous results for motor preparation have focused on hand/finger movements. For simple hand/finger movements, results have found that such movements typically evoke activity primarily in the contralateral motor cortices. However, increasing the complexity of the finger movements, such as during a distal sequential finger-pressing task, leads to additional recruitment of ipsilateral resources. It has been suggested that this involvement of the ipsilateral hemisphere is critical for temporal coordination of distal joints. The goal of the current study was to examine whether increasing simultaneous coordination of multiple joints (both proximal and distal) leads to a similar increase in coupling with ipsilateral sensorimotor cortices during motor preparation compared to a simple distal movement such as hand opening. To test this possibility, 12 healthy individuals participated in a high-density EEG experiment in which they performed either hand opening or simultaneous hand opening while lifting at the shoulder on a robotic device. We quantified within- and cross-frequency cortical coupling across the sensorimotor cortex for the two tasks using dynamic causal modeling. Both hand opening and simultaneous hand opening while lifting at the shoulder elicited coupling from secondary motor areas to primary motor cortex within the contralateral hemisphere exclusively in the beta band, as well as from ipsilateral primary motor cortex. However increasing the task complexity by combining hand opening while lifting at the shoulder also led to an increase in coupling within the ipsilateral hemisphere as well as interhemispheric coupling between hemispheres that expanded to theta, mu, and gamma frequencies. These findings demonstrate that increasing the demand of joint coordination between proximal and distal joints leads to increases in communication with the ipsilateral hemisphere as previously observed in distal sequential finger tasks.

## 1 Introduction

The majority of neuroimaging studies in humans focus on simple single-joint tasks due to practical constraints within the MRI scanner. It is clear from these studies that movements, particularly more distal ones, require communication between secondary motor regions and primary motor cortex contralateral to the moving limb^1^. Interestingly though, movements requiring greater sequential control, such as sequential finger tapping tasks, lead to increased activity in the ipsilateral sensorimotor cortex during motor preparation and execution, which appears important for temporal coordination^2–4^. However, task complexity can also be altered by changing the number of joints controlled. Considering that most activities of daily life require simultaneous coordination of multiple joints, both proximal and distal, it is important to understand how increasing the number of controlled joints may affect the reliance on the ipsilateral hemisphere.

For single-joint tasks, connectivity between motor regions has been well studied. For instance, Grefkes et al., found a facilitation of cortical activity from contralateral secondary motor areas to contralateral primary motor cortex (M1) and inhibition of ipsilateral motor areas during a simple fist closing task as measured by fMRI^1^. Similar results have been found using EEG where individuals displayed positive coupling from supplementary motor area (SMA) to contralateral motor cortices^5,6^, which matched previous findings showing activation of SMA preceding contralateral M1^7^. This evidence corroborates results from single-cell recordings in monkeys showing increased prevalence of preparation-related neurons in secondary motor areas compared to M1^8^ and suggests a cascade-like communication from secondary motor areas to M1 constrained within the contralateral hemisphere.

Although most single-joint unimanual movements are typically associated with activity in contralateral sensorimotor cortices during movement preparation and execution, the ipsilateral sensorimotor cortices seem to play a functional role as well. For instance, it is possible to decode 3D movement kinematics solely from the ipsilateral hemisphere in both monkeys^9^ and humans^10,11^, suggesting a robust role for ipsilateral sensorimotor cortices in movement preparation /execution. Meanwhile, lesioning ipsilateral M1 in monkeys leads to a brief behavioral deficit in the ipsilesional hand due to deficits in postural hand control^12^. Similarly in humans, perturbation to ipsilateral M1 via TMS leads to an increase in timing errors in tasks^4,13^, which is attributed to improper temporal recruitment of muscles^14^. Although the specific neural mechanism behind the role of the ipsilateral sensorimotor cortices in movement is still up for debate, one of the most common findings is that it plays a demand-dependent role, since increasing the ‘complexity’ of the task leads to increased activity in the ipsilateral cortex^15–17^.

Task complexity is often manipulated by having subjects execute a difficult sequence finger tapping task. However, similar results have also been found for non-sequence related tasks of increased complexity such as executing a ‘chord’ involving coordination of multiple fingers^2^. Thus, it seems that ipsilateral motor cortex is involved during preparation not only of sequential complex tasks, but also during movements that require multiple joint coordination. However, previous tasks have been limited to single-joint distal hand/finger movements or bimanual distal tasks. The question remains whether coordination of two joints within the same limb will similarly lead to increased communication with the ipsilateral hemisphere due to simultaneous control of both proximal and distal joints.

We sought to test the hypothesis that increasing coordination from a 1-joint distal task to a 2-joint distal-proximal task would increase the involvement of ipsilateral sensorimotor cortices. To investigate this hypothesis, we measured high-density EEG while participants performed either hand opening or hand opening while simultaneously lifting at the shoulder. We quantified the connectivity within bilateral sensorimotor cortices during motor preparation leading up to movement execution using dynamic causal modeling for induced responses. This allowed us to not only establish the regions involved in each task, but also disentangle the roles of different frequency coupling between tasks. We found that although both tasks displayed the expected coupling from contralateral secondary motor areas to contralateral primary motor cortex restricted to beta band, the simultaneous lifting and opening task also elicited increased coupling within the ipsilateral hemisphere towards iM1 and between contralateral PM and ipsilateral M1 that spread to theta, mu, and gamma frequencies. These results suggest that the coordination between distal and proximal joints leads to additional bilateral communication and communication within ipsilateral sensorimotor cortices compared to a simple distal movement.

## 2 Materials and Methods

### 2.1 Participants

Twelve healthy right-handed participants (mean age: 59.8 + 7.7 yrs; age range: 45-74; 7 males, 5 females) took part in this study. All participants had no prior history of neurological or psychiatric disease. This study was approved by the Northwestern institutional review board and all participants gave written informed consent.

### 2.2 Experimental Design

#### 2.2.1 Experimental setup

Participants sat in a Biodex chair (Biodex Medical Systems, Shirley, NY), with straps crossing the chest and abdomen to restrain the trunk. The participant’s right arm was placed in a forearm-hand orthosis attached to the end effector of an admittance controlled robotic device (ACT^3D^) instrumented with a six degree of freedom (DOF) load cell (JR^3^ Inc., Woodland, CA). The robot was set to the position with the height to provide a haptic table to the subject with shoulder at 85° abduction, and allows the subject move the right arm freely on the table.

At the beginning of each trial, participants moved their hand to a home position, with the shoulder at 85° abduction, 40° flexion, and the elbow at 90° flexion angle. The participant then received an auditory cue. Following the cue, participants relaxed at the home position for 5-7 s and then self-initiated either 1) hand opening (HO) with the arm resting on the haptic table, or 2) hand opening while simultaneously lifting (HOL) the arm above the haptic table against 50% of subject’s maximum shoulder abduction. Importantly, the HOL task requires simultaneous activation of the shoulder abductors and finger extensor muscles and is not a sequential task. Participants were instructed to avoid eye movements by focusing on a point and avoid movements of other body parts during the performance of each trial, which was visually confirmed by the experimenter. Participants performed 60-70 trials of each task, broken into random blocks (one block consisted of 20-30 trials for a particular task). Rest periods varied between 15 to 60 seconds between trials and 10 minutes between blocks.

#### 2.2.2 EEG Data Acquisition

Scalp recordings were made with a 160-channel High-Density EEG system using active electrodes (Biosemi, Inc, Active II, Amsterdam, The Netherlands) mounted on a stretchable fabric cap based on a 10/20 system with reflective marker on each of the electrode holders. Simultaneously, EMGs were recorded from the extensor digitorum communis (EDC), flexor carpi radialis (FCR), and intermediate deltoid (IDL) of the tested arm. All data were sampled at 2048 Hz. The impedance was kept below 50 kΩ for the duration of the experiment. The positions of EEG electrodes on the participant’s scalp were recorded with respect to a coordinate system defined by the nasion and pre-auricular notches using a Polaris Krios handheld scanner (NDI, Ontario, Canada). This allowed for coregistration of EEG electrodes with each participant’s anatomical MRI data.

#### 2.2.3 Structural Imaging of the Brain

On a different day, individuals participated in MRI scans at Northwestern University’s Center for Translation Imaging on a 3 Tesla Siemens Prisma scanner with a 64-channel head coil. Structural T1-weighted scans were acquired using an MP-RAGE sequence (TR=2.3s, TE=2.94ms, FOV 256×256mm^2^) producing an isotropic voxel resolution of 1×1×1 mm. Visual inspection of acquired images was performed immediately following the data acquisition to guarantee no artifacts and stable head position.

### 2.3 Data Analysis

#### 2.3.1 Dynamic causal modeling for induced responses

We used dynamic causal modeling for induced responses (DCM-IR)^18^ to model the task-related time-varying changes in power both within and across a range of frequencies by estimating the coupling parameters within and between sources in a network. This approach has been used in previous hand movement tasks to elucidate the dynamic frequency interactions within a motor network^5,19^.

#### 2.3.2 Definition of model space

Our network model consisted of 5 ROIs, including contralateral primary motor cortex (cM1), ipsilateral primary motor cortex (iM1), contralateral premotor cortex (cPM), ipsilateral premotor cortex (iPM), and supplementary motor area (SMA). Locations of each of these regions were adapted from the Human Motor Area Template (HMAT)^20^ and are shown in Table 1. Bilateral SMAs were treated as a single source due to their mesial position on the cortices. SMA also served as the input to the modelled network. It was chosen due to its critical role in movement preparation during self-initiated motor tasks^21,22^ and has previously been demonstrated to be an appropriate input for self-initiated motor tasks using DCM-IR^5,19,23^.

**Table 1.**
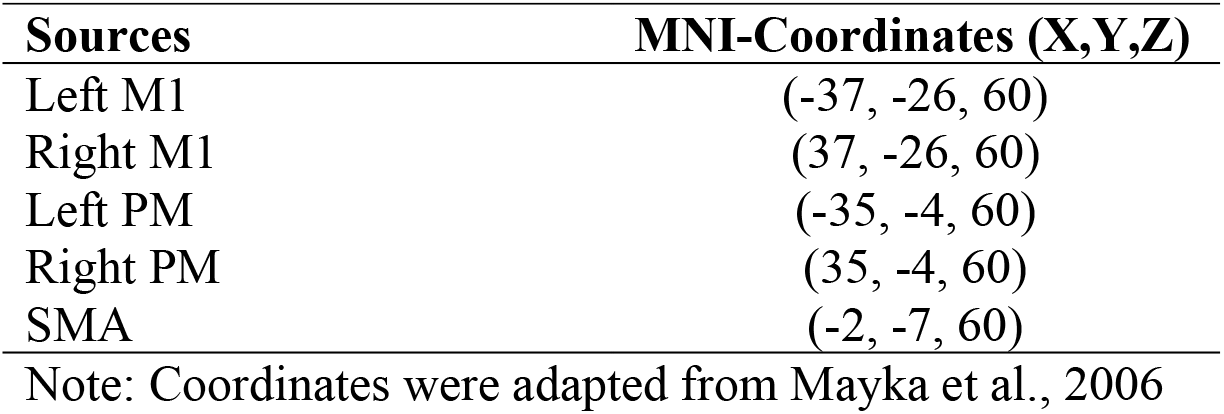
Coordinates of motor network.

| Sources | MNI-Coordinates (X,Y,Z) |
| --- | --- |
| Left M1 | (-37, -26, 60) |
| Right M1 | (37, -26, 60) |
| Left PM | (-35, -4, 60) |
| Right PM | (35, -4, 60) |
| SMA | (-2, -7, 60) |

Note: Coordinates were adapted from Mayka et al., 2006

Different within- and cross-frequency connections between these 5 sources were used to create 12 models, as shown in Supplementary Figure 1, which have successfully been used before in a grip task^19^. These 12 models were separated into 2 groups. Group 1 (models 1 to 6) allowed nonlinear and linear extrinsic (between region), but only linear intrinsic (within region) connections. Group (models 7 to 12) allowed both nonlinear and linear connections for both extrinsic and intrinsic connections. Within each group, the 6 models consisted of 1 fully connected model, and the other 5 models missing 1 or 2 connections that were from one premotor area (PM) to either the other PM or to M1. The within- and cross-frequency connections between each region from the best fit model were then used for analyzing task-related differences.

#### 2.3.3 DCM Preprocessing

EEG data were preprocessed using SPM12 (SPM12, Wellcome Trust Centre for Neuroimaging, www.fil.ion.ucl.ac.uk/spm/). Data were first band-pass filtered between 1 and 50 Hz, segmented into single trials (−2200 to 500 ms with 0 ms indicating EMG onset), and baseline-corrected. Trials were visually inspected and removed if they displayed an artifact (e.g., blinks). Artifact free trials were projected from channel space to the sources using the generalized inverse of the lead-field matrix with an equivalent current dipole for our chosen sources using a subject-specific boundary element method (BEM) based on the subject’s anatomical MRI^18^. The spectrogram of each segmented trial from 4 to 48 Hz at each source was computed using a Morlet wavelet transform. This range includes theta (4-7 Hz), mu (8-12 Hz), beta (13-35 Hz), and gamma (36-48 Hz) frequencies. The spectrogram (frequency x time x source) was then averaged over all trials, cropped between −1000 to 0 ms, and then baseline-corrected by subtracting the mean of the frequency-specific instantaneous power during the time window −1000 to −833 ms.

The dimensionality of the averaged spectrogram was then reduced to four modes (i.e., mode x time x source) using singular value decomposition (SVD). Note, after SVD we project the 45 frequencies to 4 modes. We then reshape the spectrogram to obtain the instantaneous power vector *g*20×1 at each of the sampled time, with the first 4 elements as the instantaneous power on the 1^st^ region from modes 1 to 4, and the 5^th^−8^th^ elements as the instantaneous power on the 2^nd^ region from modes 1-4, and so on. This dimensionality reduction both reduced the computational demand of the model inversion and denoised the data.

#### 2.3.4 Calculation of coupling parameters

Using the models shown in Supplementary Fig 1, we simulated dynamics of the instantaneous power using the following equation:

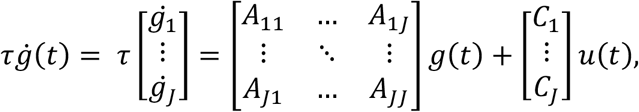

Where the vector 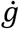 represents the first derivative of instantaneous power *g*. The sub-matrix *A_ij_* is a 4 by 4 matrix containing the coupling parameters within and across different modes between the *i*^*th*^ and *j*^*th*^ regions (*J* = 5). The vector *u* represents the extrinsic input, which, in this study, is modeled as a gamma function with a peak at 400 ms prior to EMG onset with a dispersion of 400 ms from SMA to the whole network. These values were chosen in order to capture the peak of the bereitschaftspotential during a self-initiated movement^24^. The C matrix contains the weights of the extrinsic input *u* from SMA. τ is a scaling factor and *t* represents time.

The model simulation was restricted to the time leading up to EMG onset (−1000 to 0 ms) to capture purely the motor preparation and command rather than any potential sensory feedback related to the task. We optimized the A and C matrices to minimize the error between the measured and simulated spectrogram. The quality of a model and the estimated A and C matrices was quantified by the variance accounted for from the simulated spectrogram. The resulted elements in the coupling A matrix refers to the influence of power at a specific frequency in one motor region on the power at another frequency in another region (column to row). Positive (i.e., excitatory) or negative (i.e., inhibitory) coupling suggests changes in power in the first frequency and region lead to the same directional or opposite change, respectively, in power in the second frequency and region.

#### 2.3.5 Bayesian model selection

We performed Bayesian model selection (BMS) with random effects to assess which model best explained the observed data, while taking into account the complexity of the model^25,26^. We first used a family level inference with random effects to assess the overall importance of nonlinear coupling in intrinsic (i.e., within-region) connections. This involved comparing models 1-6 (Linear intrinsic connections) with models 7-12 (Linear and Nonlinear intrinsic connections). Following evaluation at the family level, we used BMS on the 6 models from the winning family to see which model best explained the observed data. The winning model, which was then used for further analysis on task-related differences in coupling, was chosen based on the highest posterior exceedance probability (i.e., the probability that a given model is more likely than any of the other models tested). This process was performed for both tasks separately.

#### 2.3.6 Inference on coupling parameters

Simulated spectrograms and A matrices from the four modes were projected back to frequency domain allowing for characterization of the coupling parameters as a function of frequency for the winning model. The coupling matrices for each intrinsic and extrinsic connections in the winning model for each participant were further smoothed with a Gaussian kernel (full-width half-maximum of 8 Hz) for each condition. These matrices include the frequency-to-frequency (both within- and cross-frequency) coupling values for each connection. We then ran a one-sample *t*-test for the coupling parameters for each connection to assess connections involved in the default network for each condition. We then ran a paired *t-*test on the coupling parameters on significant connections of the default network to assess task-related differences in coupling. Significance for specific coupling parameters was considered statistically significant at *p* < 0.005 uncorrected.

## 3 Results

### 3.1 Behavioral Results

We first sought to confirm that participants were simultaneously activating the hand and shoulder during the HOL task. We found that participants were reliably able to activate both muscles simultaneously, showing an absolute difference of EMG activation between muscles of 75.7 ± 54.6 ms (median = 57.6 ms; min = 36.5 ms; max = 161.0 ms) on average (see Figure 1). A histogram of this absolute difference of EMG activation between muscles for all trials and participants is shown in Figure 1B.

**Figure 1.**
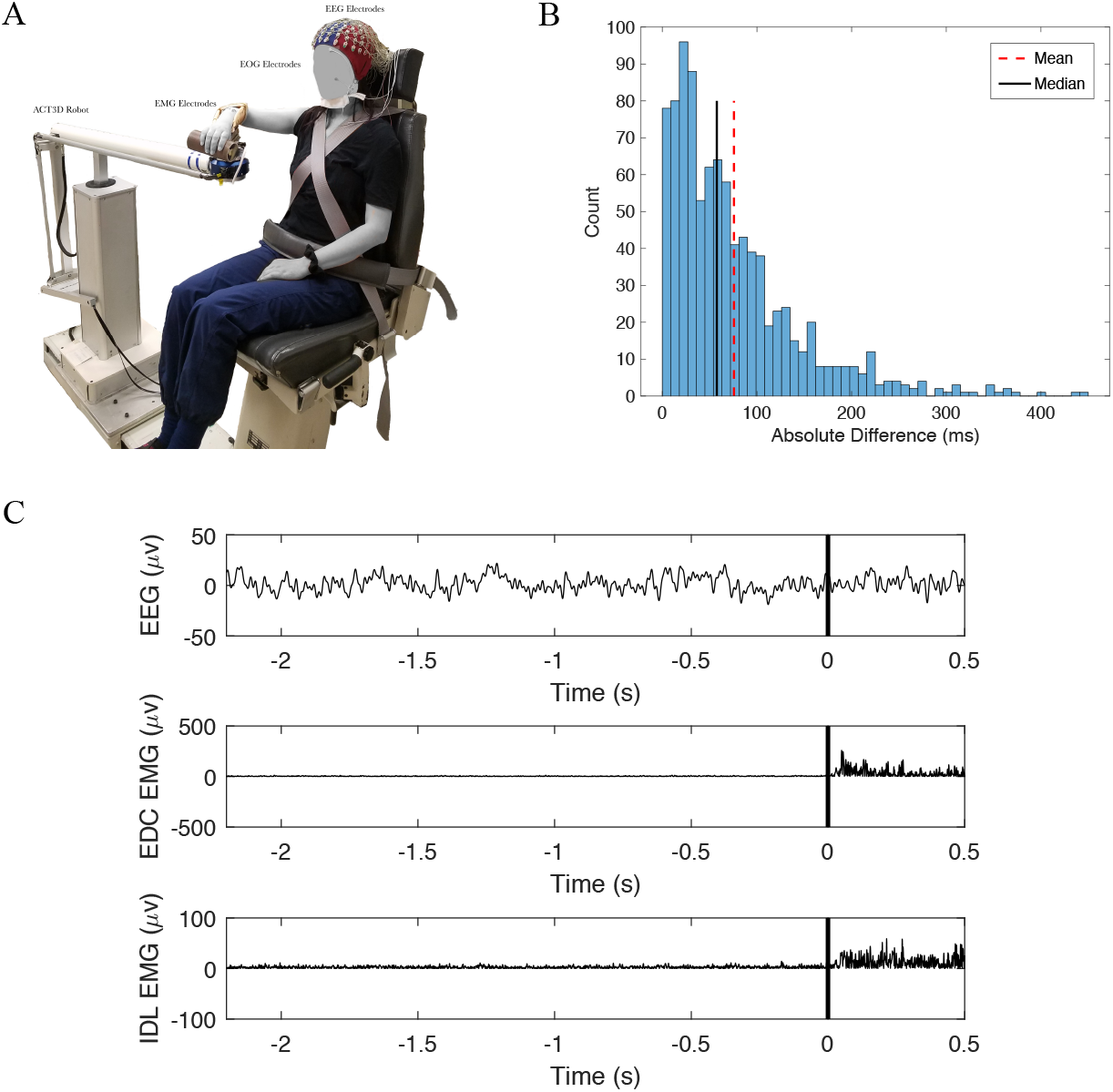
(A) EEG setup on the ACT3D robot. (B) Histogram of the absolute difference in onset between the extensor digitorum communis (EDC) and the deltoid (IDL) across all trials and participants. The mean absolute difference is represented by the red dashed vertical line and the median is represented by the solid black vertical line. (C) Example from one trial of EEG data (top) and extensor (middle) and deltoid (bottom) EMG signal. Black vertical line at 0 s represents EMG onset across all 3 signals.

### 3.2 Bayesian model selection and model fit

Results of BMS were consistent across the two tasks. Family-level inference showed the strongest evidence for the Nonlinear family, which allowed both within- and cross-frequency coupling for intrinsic (i.e., within-region) and extrinsic (i.e., between-region) connections (Exceedance probabilities for HO: 0.9999 and HOL: 0.9998; Supplementary Table 1). When comparing the six models from the Nonlinear family (models 7-12 in Supplementary Figure 1), BMS favored Model 12, which contained full connections between the 5 regions of interest (Exceedance probabilities for HO: 0.9994 and HOL: 0.9981; Supplementary Table 2).

Supplementary Figure 2 depicts both the observed and simulated spectrograms in each of the 5 motor regions using the winning model for one participant during HO. Comparison of these two spectrograms shows the overall similarity between the measured) and the model-based simulated data. Overall, this model explained ~80% of the original spectral variance for each condition (HO: 82.0%; HOL: 79.3%). Additionally, the four modes from the SVD preserved >95% of the data variance on average (HO: 95.3%; HOL: 95.7%).

Figure 2 shows the simulated spectrograms for each of the 5 motor regions using the winning model for both HO (Figure 2A) and HOL (Figure 2B). A strong β band (13-35 Hz) desynchronization is observed in the time leading up to movement onset, particularly in cM1, iM1, cPM, and SMA for both tasks. Additionally, cM1, cPM, and SMA show γ band (36-48 Hz) synchronization just before movement onset (starting ~100 ms before EMG onset).

**Figure 2.**
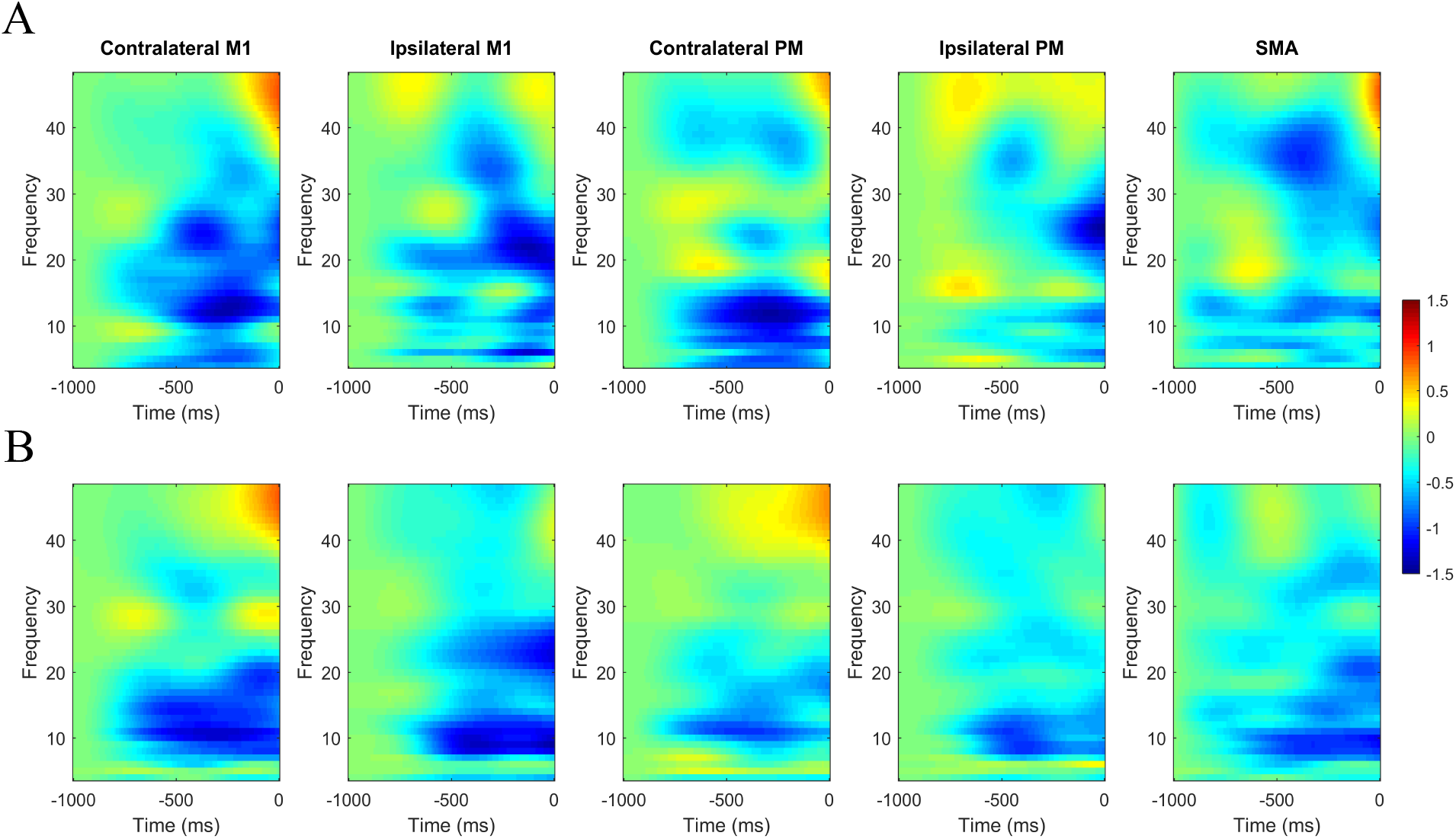
Time-Frequency plot for (A) Hand Opening (HO) and (B) Hand Opening while Lifting (HOL) for the 5 regions of interest. Blue depicts decreases in power relative to baseline and red depicts increases in power relative to baseline. 0 ms indicates EMG onset.

### 3.3 Default motor networks for the two tasks

We then evaluated the default motor networks for the two tasks. The HO task demonstrated significant positive (i.e., excitatory) coupling from contralateral secondary motor areas to contralateral primary motor cortex. This included coupling from SMA to cM1, cPM to cM1, and SMA to cPM (see Figure 3A), all confined to the beta band (13-35 Hz). Additionally, individuals displayed positive interhemispheric coupling from iM1 to cM1 within the beta band.

**Figure 3.**
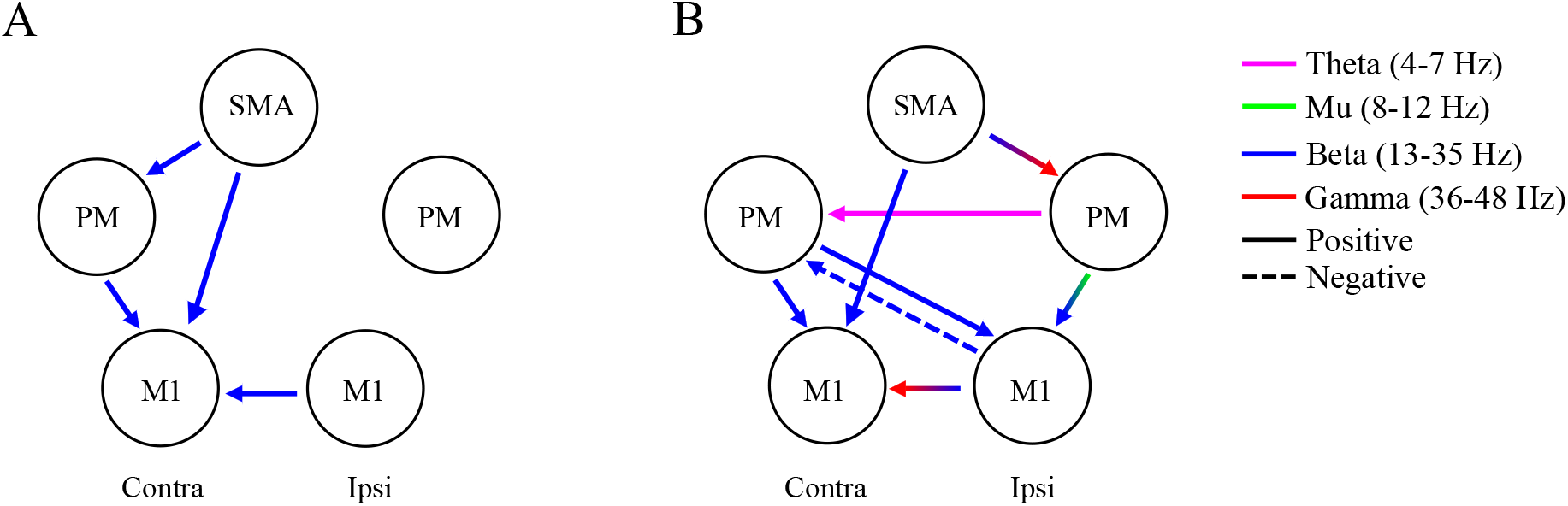
Default Oscillatory Coupling for (A) Hand Opening (HO) and (B) Hand Opening while Lifting (HOL). Arrows indicate directional connections showing significant coupling within the motor network. The color of the arrow indicates the frequency band involved. Arrows that change colors represent cross-frequency coupling. Solid lines indicate positive coupling while dashed lines indicate negative coupling. Contra = Contralateral hemisphere; Ipsi = Ipsilateral hemisphere.

During the HOL task, individuals also demonstrated significant positive coupling from contralateral secondary motor areas to contralateral primary cortex. This included coupling from SMA to cM1 and from cPM to cM1 (see Figure 3B), again all confined to beta band. However, in addition to this coupling, individuals displayed more complex ipsilateral and cross hemisphere coupling. This included positive interhemispheric coupling from iM1 to cM1 once again but involving beta and now gamma (36-48 Hz) oscillations. Additionally, HOL showed coupling (positive and negative) both within the ipsilateral hemisphere and across hemispheres (see Figure 3B). These connections spread across multiple frequency bands, including theta (4-7 Hz), alpha (8-12 Hz), beta and gamma frequencies (36-48 Hz). Table 2 contains the full characteristics of each significant connection for the two tasks.

**Table 2.** Significant default frequency-to-frequency coupling.

| Connection | Frequency Bands<br>(peak [Hz]) | # of Voxels | T-Value | Excitatory/Inhibitory |
| --- | --- | --- | --- | --- |
| **Condition:**<br>**Open** |  |  |  |  |
| SMA → cM1 | $\beta \rightarrow \beta$ (25 → 24) | 14 | 2.7 | + |
| cPM → cM1 | $\beta \rightarrow \beta$ (27 → 20) | 10 | 3.0 | + |
| SMA → cPM | $\beta \rightarrow \beta$ (15 → 13) | 60 | 4.0 | + |
| iM1 → cM1 | $\beta \rightarrow \beta$ (15 → 32) | 15 | 3.6 | + |
| **Condition:**<br>**Open while**<br>**Lifting** |  |  |  |  |
| SMA → cM1 | $\beta \rightarrow \beta$ (34 → 19) | 11 | 3.5 | + |
| cPM → cM1 | $\beta \rightarrow \beta$ (26 → 23) | 16 | 3.4 | + |
| iM1 → cM1 | $\beta \rightarrow \gamma$ (24 → 39) | 34 | 3.4 | + |
| cPM → iM1 | $\beta \rightarrow \beta$ (30 → 17) | 40 | 3.8 | + |
| | $\alpha \rightarrow \beta$ (12 → 35) | 36 | 4.6 | + |
| iM1 → cPM | $\beta \rightarrow \beta$ (33 → 25) | 102 | 4.9 | - |
| iPM → cPM | $\phi \rightarrow \phi$ (7 → 7) | 14 | 3.8 | + |
| SMA → iPM | $\beta \rightarrow \gamma$ (18 → 36) | 8 | 3.4 | + |
| iPM → iM1 | $\mu \rightarrow \beta$ (11 → 14) | 31 | 3.7 | + |
| | $\beta \rightarrow \beta$ (32 → 13) | 8 | 3.3 | + |

Notes: # of voxels refers to the spread of coupling around the peak frequency involved where each voxel represents coupling with 1 Hz resolution; Excitatory refers to positive coupling where a change in power in the 1<sup>st</sup> connection leads to the same directional change in power in the 2<sup>nd</sup> connection; Inhibitory refers to negative coupling where a change in power in the 1<sup>st</sup> connection leads to the opposite directional change in power in the 2<sup>nd</sup> connection; Highlighted rows reflect common connections in both tasks. M1 = Primary Motor Cortex; PM = Premotor Cortex; SMA = Supplementary Motor Area; c = contralateral; i = Ipsilateral

### 3.4 Task-related differences in coupling

We observed significant differences in coupling between the two tasks. Overall, the cortical network for preparing the HOL task was more complex than that for HO task, as apparent from its increased involvement of additional network from ipsilateral motor cortices. This included from SMA to iPM and iPM to iM1 across multiple frequency bands (theta, beta, and gamma; see Figure 4) compared to HO. Additionally, the HOL task showed greater positive interhemispheric coupling from cPM to iM1 and more negative coupling from iM1 to cPM within the beta band compared to HO. The only significantly stronger link for the HO task is from iM1 to cM1 within the beta band compared to HOL. Table 3 contains the full characteristics of each significant connection for the two-task comparison.

**Table 3.**
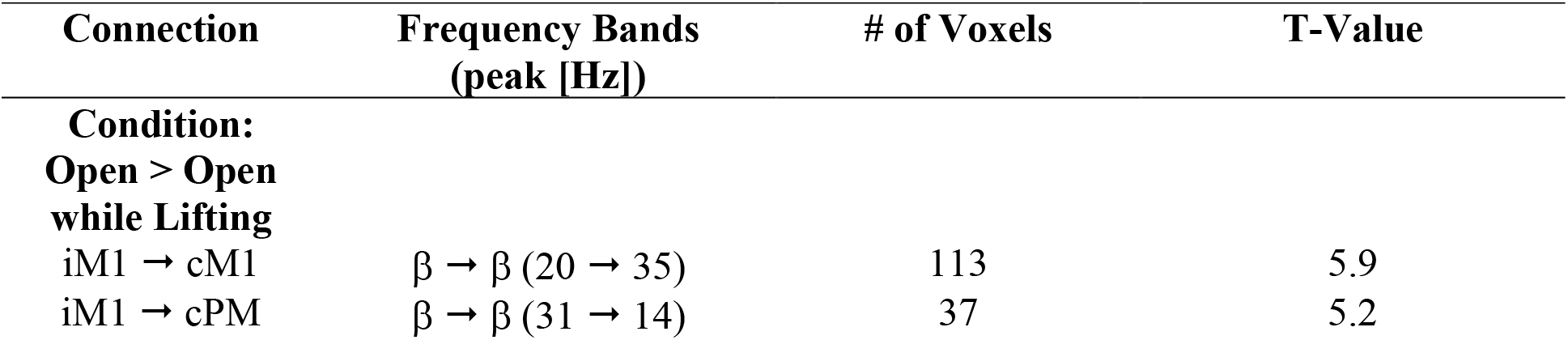
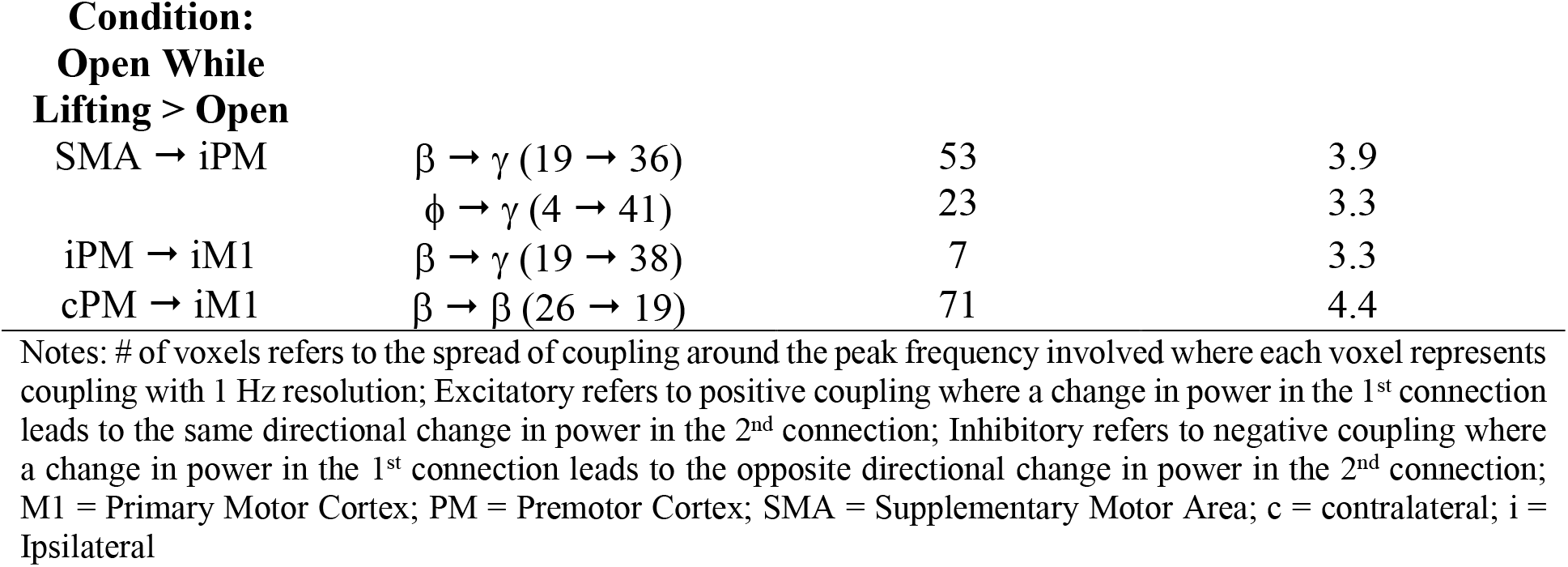
Significant differences in frequency-to-frequency coupling between tasks.

| Connection | Frequency Bands<br>(peak [Hz]) | # of Voxels | T-Value |
| --- | --- | --- | --- |
| **Condition:**<br>**Open > Open**<br>**while Lifting** |  |  |  |
| iM1 → cM1 | $\beta \rightarrow \beta$ (20 → 35) | 113 | 5.9 |
| iM1 → cPM | $\beta \rightarrow \beta$ (31 → 14) | 37 | 5.2 |
| **Condition: Open While Lifting > Open** |  |  |  |
| SMA → iPM | $\beta \rightarrow \gamma$ (19 → 36) | 53 | 3.9 |
| | $\phi \rightarrow \gamma$ (4 → 41) | 23 | 3.3 |
| iPM → iM1 | $\beta \rightarrow \gamma$ (19 → 38) | 7 | 3.3 |
| cPM → iM1 | $\beta \rightarrow \beta$ (26 → 19) | 71 | 4.4 |

Notes: # of voxels refers to the spread of coupling around the peak frequency involved where each voxel represents coupling with 1 Hz resolution; Excitatory refers to positive coupling where a change in power in the 1<sup>st</sup> connection leads to the same directional change in power in the 2<sup>nd</sup> connection; Inhibitory refers to negative coupling where a change in power in the 1<sup>st</sup> connection leads to the opposite directional change in power in the 2<sup>nd</sup> connection; M1 = Primary Motor Cortex; PM = Premotor Cortex; SMA = Supplementary Motor Area; c = contralateral; i = Ipsilateral

**Figure 4.**
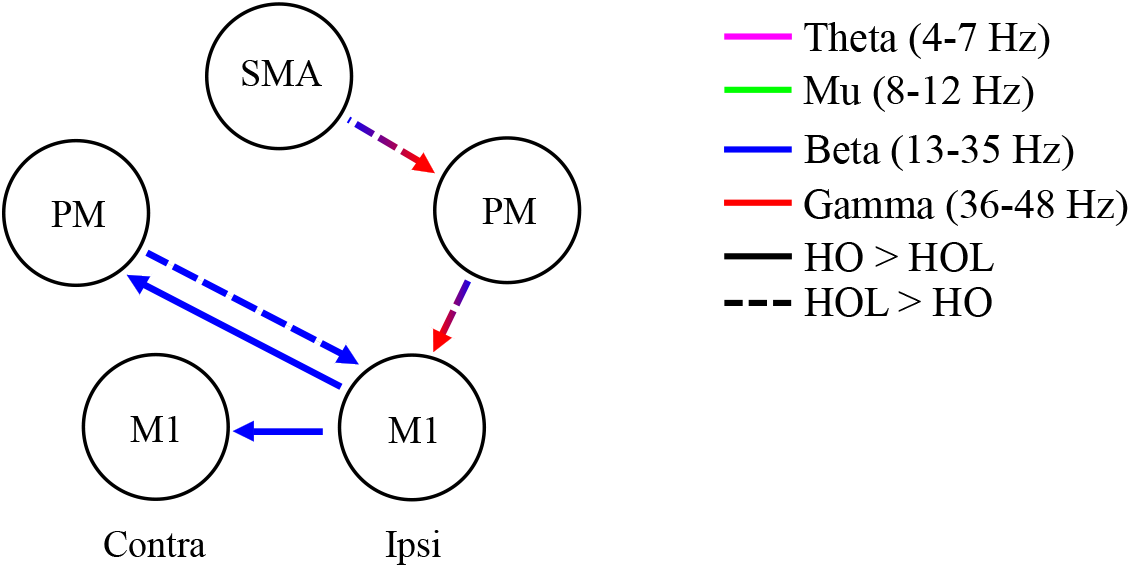
Differences in oscillatory coupling between the two tasks. Arrows indicate directional connections showing significant differences in coupling between tasks within the motor network. The color of the arrow indicates the frequency band involved. Arrows that change colors represent cross-frequency coupling. Solid lines indicate greater coupling for Hand Opening (HO) compared to Hand Opening while Lifting (HOL), while dashed lines indicate greater coupling for HOL compared to HO.

## 4 Discussion

This paper sought to investigate whether increasing coordination from a pure hand opening (HO) task to a simultaneous hand opening while lifting (HOL) task that requires coordination between the shoulder and hand would increase involvement of the ipsilateral hemisphere. For the first time, we established the default network (i.e., cortical-cortical coupling at different frequencies) in the time leading up to movement execution for these two movements. Then we compared the difference between these default networks for the two tasks and showed that 1) contralateral beta-to-beta coupling between SMA/cPM and cM1 is commonly involved in both the HO and HOL tasks; and 2) increased bilateral connectivity and connectivity within the ipsilateral sensorimotor cortices across multiple frequency bands are involved in the HOL task, a task requiring coordination between the shoulder and hand, but not in the HO task.

### 4.1 Common Networks During Single- and Multi-Joint Movements

#### 4.1.1 Common Network

Both tasks evoked a common excitatory coupling pattern from secondary motor cortices (SMA and cPM) to motor cortex (cM1) in the contralateral hemisphere in the beta frequency band (13-35 Hz). The involvement of secondary motor cortices falls in line with previous proposed roles for these regions: secondary motor areas feed into primary motor cortex during motor preparation and execution to shape motor output^27–31^. Such increased preparatory activity from contralateral secondary motor areas was observed in both distal hand tasks^32^ as well as multi-joint movements involving the shoulder, such as reaching^33^. In regard to the frequencies involved, the positive beta coupling reflects the common beta desynchronization seen across these regions in the time-frequency maps (see Figure 2), which is a feature of movement preparation in both secondary and primary motor cortices. This beta band desynchronization is associated with the gradual release of inhibition in the motor cortex to initiate an action^34,35^, along with the descending motor command originating from layer V pyramidal cells^36,37^. We posit that the presence of beta band coupling within secondary and primary motor areas in the contralateral hemisphere observed for the two tasks in this experiment reflects this common motor command for the two tasks.

### 4.2 Distinct Networks During Single- and Multi-Joint Movements

We provided evidence, for the first time, showing the increased involvement of the ipsilateral hemisphere during the preparation of the simultaneous lifting and hand opening (HOL) task, as compared to the pure hand opening (HO) task. Specifically, the HOL task elicited coupling within the ipsilateral hemisphere (SMA to iPM; iPM to iM1), as well as between the 2 hemispheres (i.e., bidirectional link between iM1 and cPM) during movement preparation.

#### 4.2.1 Significant Differences between the 2 tasks

One of the significant between-task differences in the network was the presence of ipsilateral beta-to-gamma coupling both from SMA to iPM and from iPM to iM1 during HOL task. As beta oscillations are an index of inhibition^38^, beta ERD, as seen in SMA at −450 ms at 19 Hz and iPM around at −500 ms during HOL task in Figure 2, may suggest that the reduction of such inhibition facilitated the synchronization of targeted regions (i.e., iPM and iM1) at gamma band. Neural firing at gamma oscillations (30-80 Hz) on the superficial layers of the frontal cortex is believed to be generated by the loop of inhibition between fast spiking GABAergic interneurons and pyramidal cells^39^. Therefore, the resulting gamma hyper-synchronization in iPM and iM1 may represent a shift in connectivity away from long-range interlaminar connectivity typically associated with slower frequencies such as beta towards more local circuits^40^. The computational results within the local circuits may further drive cells from deeper cortical layers, as demonstrated by the fact that gamma oscillations can drive the connections to target cells in both superficial and deeper cortical layers^41^. This is different from the commonly involved beta-to-beta coupling at the contralateral side also from the secondary to the primary motor cortex, which may purely reflect the release of the prepared motor plan without the local computation that primarily occurs here in the ipsilateral motor cortices at higher frequencies.

The other significant distinct coupling for the HOL task is a bidirectional beta band coupling between iM1 and cPM, with cPM synchronization facilitating iM1 synchronization, whereas iM1 synchronization inhibits cPM synchronization. An anatomical link between iM1 and cPM has been reported via corpus callosum^42^, although it is also possible this may reflect communication through hidden or additional nodes (either subcortical or cortical) not included in our motor network. In Figure 2, we observed increased synchronization in cPM around at 30 Hz, which was coupled with the increased synchronization in iM1 around at 19 Hz. This may reflect increased inhibition of iM1 activity initiated by cPM inhibition. On the other hand, the beta synchronization at high beta in iM1 increased and triggered beta desynchronization in cPM at low beta (~14 Hz) component, suggesting a release of inhibition to cPM activity. Overall, this loop may result in an increase in cortical activity at contralateral secondary motor cortices within the beta band, and an inhibition of activity in ipsilateral primary motor cortex.

The last significant difference in coupling between the 2 tasks was the shift of facilitation coupling from iM1 to cM1 in the beta band for HO task to a beta-to-gamma coupling still from iM1 to cM1 for HOL task. Ipsilateral M1 is known to communicate with cM1 via transcallosal connections starting during movement preparation through movement onset^42,43^, and thus only the nature of this coupling (i.e., frequencies involved) seem to be changing based on the particular task. One possibility for the larger beta-beta coupling for the HO task is due to the distal-only nature of the task compared to the proximal-distal combination for the HOL task, as previous findings showed greater beta event-related desynchronization contralateral to the moving limb for distal finger movements compared to proximal shoulder movements^44^.

#### 4.2.2 Other differences between the 2 tasks

When comparing the within and cross-frequency couplings between the two tasks listed in table 2, there are several more couplings that were involved in HOL task, but not in the HO task, although further paired t-test did not report them as significant difference. One of them is the facilitative theta-to-theta coupling from iPM to cPM. Theta (4-7 Hz) oscillation have been implicated in long range integration and top-down processing for various tasks in the cognitive domain, showing higher power with increasing cognitive demand^45–47^. Considering that the simultaneous lifting and opening task did require increased coordination of multiple joints compared to the simple hand opening task, coupling between iPM and cPM in theta band might reflect the increased cognitive load for the HOL task as compared to HO task. The fact that theta coupling was prevalent only in the premotor areas rather than primary motor cortex further implies that this theta coupling may be more indicative of cognitive processes rather than purely a motor command, as premotor areas are typically associated with more abstract and goal-directed representations of movement compared to M1^48^.

We also observed mu-to-beta coupling, uniquely in HOL task, between cPM/iPM and iM1. Here, we have referred to this band as mu since it is part of the alpha band related to movement. As shown in Figure 2, we observed a strong mu-wave suppression (at about −550 ms) in both iPM and cPM leading to the beta suppression (at about −500 ms) in iM1 for HOL task but not HO task. Suppression of mu waves are commonly observed when one performs or visualizes performing a motor action. Oscillations at these lower frequencies may be more present in deeper cortical layers, which then innervate superficial cortical layers^49,50^. The slower firing properties of deeper layers are more appropriate to synchronize cell assemblies over longer conduction delays^40^. Based on these previous results, it is possible that the observed mu to beta coupling from bilateral PMs to iM1 may indicate the use of deeper structures for the long-scale cross-hemisphere communication to iM1. This may be necessary for the higher level of coordination which is required for multi-joint task like HOL. In line with this notion, previous findings showed that elderly individuals displayed greater spread of mu-suppression across primary and secondary motor areas during a self-paced thumb movement compared to younger individuals, probably due to having to put more effort into the task^51^. Another possibility is that this observed mu-beta coupling reflects use of descending motor pathways controlling the shoulder, as ipsilateral mu suppression has been linked with excitability of uncrossed pathways projecting to shoulder muscles^52^.

### 4.3 Potential Role of the Ipsilateral Hemisphere

Since increasingly difficult finger tasks elicit increased activity in the ipsilateral hemisphere^2,3^, we hypothesized and confirmed that the HOL task would similarly increase connectivity with the ipsilateral hemisphere compared to the HO task due to the increased complexity of simultaneously coordinating proximal and distal joints. However, the question remains what the overall potential role of the ipsilateral hemisphere involvement may be for the HOL task.

The observed ipsilateral connectivity for the HOL task may suggest that ipsilateral motor cortices are directly involved in the preparation and/or execution of the more complex movement. In line with this possibility, Horenstein and colleagues compared the amount of cortical activity in the ipsilateral sensorimotor cortex during a unimanual and bimanual complex finger tapping task^53^. They found a large amount of overlap of activity in the ipsilateral hemisphere during the two tasks. Since the activity in the ipsilateral hemisphere during movement of the ipsilateral hand overlapped with the activity during bimanual movement, they argued the overlapping activity was presumably due to the preparation and execution of the movement itself. This evidence fits well with previous decoding studies showing the ability to decode 3D movements purely from activity in the ipsilateral hemisphere^10,11^ and that these ipsilateral representations seem to be related to active movement rather than sensory processes (as is likely the case here as well since analyses were restricted to the time leading up to EMG onset)^54^.

Considering the HOL task required simultaneous coordination of both the hand and the shoulder, it is also possible that the ipsilateral hemisphere plays a role in synchronizing the timing of recruitment of the muscles involved in the movement via transcallosal mechanisms. In support of this potential role, virtual lesions to ipsilateral M1 elicited by TMS have been shown to alter the timing of muscle recruitment and lead to significant motor deficits during a multi-joint grip-lift task^14^. Similarly, inhibitory TMS over ipsilateral M1 led to temporal alterations in the sequence of finger tapping movements of increasing complexity, but without affecting the number of incorrect sequences of the movements^13^. Therefore, it is possible that the increased coupling with the ipsilateral hemisphere observed here plays a significant role in coordinating the simultaneous activation of both proximal and distal joints. The observed role for gamma coupling may facilitate this due to its role in local computation and GABAergic inhibitory circuity^40,55^. Meanwhile, an alternative possibility that others have suggested is that this ipsilateral activity reflects inhibition of possible mirror movements of the ipsilateral hand rather than just an interhemispheric control mechanism^56^.

### 4.4 Limitations

We cannot fully rule out the possibility that the observed increase in ipsilateral connectivity during the multi-joint task indicates recruitment of descending uncrossed motor tracts from the ipsilateral hemisphere such as the ipsilateral corticospinal tract or cortico-reticulospinal tract. Although this is unlikely for the sequential finger tasks due a lack of innervation of these pathways to distal portions of the hand^57^, it is potentially relevant for the task in this study as these pathways have been shown to have substantial connections to more proximal portions of the upper extremity, such as the deltoid that is involved in the lifting portion of the task^58^. Therefore, the observed increase in ipsilateral connectivity may reflect recruitment of ipsilateral descending motor tracts to drive the shoulder during the lifting task, with the contralateral hemisphere still providing the majority of the input for controlling the distal hand opening.

Furthermore, due to the lack of a lifting-only condition, i.e., a task only involving shoulder joint, it is possible that a portion of the task-related changes in connectivity are solely related to the lifting component of the movement rather than the combination of simultaneously opening the hand and lifting at the shoulder. However, previous evidence has shown that activity during the motor preparation phase of a lifting-only single joint movement is primarily restricted to contralateral motor cortex and secondary motor areas with minimal ipsilateral involvement^59^.

Other limitations of the presented study are associated with the use of the DCM-IR method. This method only takes into account the temporal changes in power of particular frequencies but does not account for phase. Phase is known to play a critical role in cognitive and sensorimotor processes separate from power/amplitude^60^. Another limitation is that DCM-IR is limited in the number of sources that can be included in the model. However, we believe the tasks and ROIs chosen in this study are well-justified by previous work, and although they certainly do not characterize the entirety of the network involved in these tasks, we believe they carry enough information to make worthwhile conclusions about the impact of increasing task-complexity via simultaneous control of multiple joints on cortical communication.

## 5 Conclusion

The current study demonstrated that increasing task-complexity from controlling one joint (i.e., hand opening) to coordinating multiple joints simultaneously (i.e., hand opening while lifting) led to an increase in coupling within the ipsilateral sensorimotor cortex and between hemispheres. Different from the common beta-to-beta coupling in the contralateral hemisphere, ipsilateral coupling involves a wide range of within- and cross-frequency coupling including theta, mu, and gamma frequencies. These results suggest that complexity-related reliance on the ipsilateral hemisphere holds true not just for complex sequential finger tasks, but also during combined distal-proximal multi-joint tasks more relevant to many activities of daily life.

## Acknowledgements

The authors want to acknowledge Dylan Fitzsimons and Carolina Carmona for help with data collection.

## Ethical Approval

All procedures performed in studies involving human participants were in accordance with the ethical standards of the institutional and/or national research committee with the 1964 Helsinki declaration and its later amendments or comparable ethical standards.

## References

1. Grefkes C, Eickhoff SB, Nowak DA, Dafotakis M, Fink GR. Dynamic intra- and interhemispheric interactions during unilateral and bilateral hand movements assessed with fMRI and DCM. NeuroImage. 2008;41(4):1382–1394.

2. Verstynen T, Diedrichsen J, Albert N, Aparicio P, Ivry RB. Ipsilateral motor cortex activity during unimanual hand movements relates to task complexity. Journal of neurophysiology. 2005;93(3):1209–1222.

3. Tanji J, Okano K, Sato KC. Neuronal activity in cortical motor areas related to ipsilateral, contralateral, and bilateral digit movements of the monkey. Journal of neurophysiology. 1988;60(1):325–343.

4. Chen R, Gerloff C, Hallett M, Cohen LG. Involvement of the ipsilateral motor cortex in finger movements of different complexities. Annals of neurology. 1997;41(2):247–254.

5. Bonstrup M, Schulz R, Feldheim J, Hummel FC, Gerloff C. Dynamic causal modelling of EEG and fMRI to characterize network architectures in a simple motor task. NeuroImage. 2015;124(Pt A):498–508.

6. Herz DM, Christensen MS, Reck C, et al. Task-specific modulation of effective connectivity during two simple unimanual motor tasks: a 122-channel EEG study. NeuroImage. 2012;59(4):3187–3193.

7. Huang MX, Harrington DL, Paulson KM, Weisend MP, Lee RR. Temporal dynamics of ipsilateral and contralateral motor activity during voluntary finger movement. Human brain mapping. 2004;23(1):26–39.

8. Riehle A, Requin J. Monkey primary motor and premotor cortex: single-cell activity related to prior information about direction and extent of an intended movement. Journal of neurophysiology. 1989;61(3):534–549.

9. Ganguly K, Secundo L, Ranade G, et al. Cortical representation of ipsilateral arm movements in monkey and man. The Journal of neuroscience : the official journal of the Society for Neuroscience. 2009;29(41):12948–12956.

10. Bundy DT, Szrama N, Pahwa M, Leuthardt EC. Unilateral, 3D Arm Movement Kinematics Are Encoded in Ipsilateral Human Cortex. The Journal of neuroscience : the official journal of the Society for Neuroscience. 2018;38(47):10042–10056.

11. Hotson G, Fifer MS, Acharya S, et al. Coarse electrocorticographic decoding of ipsilateral reach in patients with brain lesions. PloS one. 2014;9(12):e115236.

12. Bashir S, Kaeser M, Wyss A, et al. Short-term effects of unilateral lesion of the primary motor cortex (M1) on ipsilesional hand dexterity in adult macaque monkeys. Brain Struct Funct. 2012;217(1):63–79.

13. Avanzino L, Bove M, Trompetto C, Tacchino A, Ogliastro C, Abbruzzese G. 1-Hz repetitive TMS over ipsilateral motor cortex influences the performance of sequential finger movements of different complexity. The European journal of neuroscience. 2008;27(5):1285–1291.

14. Davare M, Duque J, Vandermeeren Y, Thonnard JL, Olivier E. Role of the ipsilateral primary motor cortex in controlling the timing of hand muscle recruitment. Cerebral cortex. 2007;17(2):353–362.

15. Buetefisch CM, Revill KP, Shuster L, Hines B, Parsons M. Motor demand-dependent activation of ipsilateral motor cortex. Journal of neurophysiology. 2014;112(4):999–1009.

16. Hummel F, Kirsammer R, Gerloff C. Ipsilateral cortical activation during finger sequences of increasing complexity: representation of movement difficulty or memory load? Clinical neurophysiology : official journal of the International Federation of Clinical Neurophysiology. 2003;114(4):605–613.

17. Seidler RD, Noll DC, Thiers G. Feedforward and feedback processes in motor control. NeuroImage. 2004;22(4):1775–1783.

18. Chen CC, Kiebel SJ, Friston KJ. Dynamic causal modelling of induced responses. NeuroImage. 2008;41(4):1293–1312.

19. Chen CC, Kilner JM, Friston KJ, Kiebel SJ, Jolly RK, Ward NS. Nonlinear coupling in the human motor system. The Journal of neuroscience : the official journal of the Society for Neuroscience. 2010;30(25):8393–8399.

20. Mayka MA, Corcos DM, Leurgans SE, Vaillancourt DE. Three-dimensional locations and boundaries of motor and premotor cortices as defined by functional brain imaging: a meta-analysis. NeuroImage. 2006;31(4):1453–1474.

21. Jahanshahi M, Jenkins IH, Brown RG, Marsden CD, Passingham RE, Brooks DJ. Self-initiated versus externally triggered movements. I. An investigation using measurement of regional cerebral blood flow with PET and movement-related potentials in normal and Parkinson's disease subjects. Brain : a journal of neurology. 1995;118 (Pt 4):913–933.

22. Jenkins IH, Jahanshahi M, Jueptner M, Passingham RE, Brooks DJ. Self-initiated versus externally triggered movements. II. The effect of movement predictability on regional cerebral blood flow. Brain : a journal of neurology. 2000;123 (Pt 6):1216–1228.

23. Loehrer PA, Nettersheim FS, Jung F, et al. Ageing changes effective connectivity of motor networks during bimanual finger coordination. NeuroImage. 2016;143:325–342.

24. Shibasaki H, Hallett M. What is the Bereitschaftspotential? Clinical neurophysiology : official journal of the International Federation of Clinical Neurophysiology. 2006;117(11):2341–2356.

25. Penny WD, Stephan KE, Mechelli A, Friston KJ. Comparing dynamic causal models. NeuroImage. 2004;22(3):1157–1172.

26. Stephan KE, Penny WD, Daunizeau J, Moran RJ, Friston KJ. Bayesian model selection for group studies. NeuroImage. 2009;46(4):1004–1017.

27. Lara AH, Cunningham JP, Churchland MM. Different population dynamics in the supplementary motor area and motor cortex during reaching. Nature communications. 2018;9(1):2754.

28. Ohara S, Ikeda A, Kunieda T, et al. Movement-related change of electrocorticographic activity in human supplementary motor area proper. Brain : a journal of neurology. 2000;123 (Pt 6):1203–1215.

29. Sun H, Blakely TM, Darvas F, et al. Sequential activation of premotor, primary somatosensory and primary motor areas in humans during cued finger movements. Clinical neurophysiology : official journal of the International Federation of Clinical Neurophysiology. 2015;126(11):2150–2161.

30. Weilke F, Spiegel S, Boecker H, et al. Time-resolved fMRI of activation patterns in M1 and SMA during complex voluntary movement. Journal of neurophysiology. 2001;85(5):1858–1863.

31. Chen CF, Kreutz-Delgado K, Sereno MI, Huang RS. Unraveling the spatiotemporal brain dynamics during a simulated reach-to-eat task. NeuroImage. 2019;185:58–71.

32. Okano K, Tanji J. Neuronal activities in the primate motor fields of the agranular frontal cortex preceding visually triggered and self-paced movement. Experimental brain research. 1987;66(1):155–166.

33. Picard N, Strick PL. Activation of the supplementary motor area (SMA) during performance of visually guided movements. Cerebral cortex. 2003;13(9):977–986.

34. Takemi M, Masakado Y, Liu M, Ushiba J. Sensorimotor event-related desynchronization represents the excitability of human spinal motoneurons. Neuroscience. 2015;297:58–67.

35. Takemi M, Masakado Y, Liu M, Ushiba J. Event-related desynchronization reflects downregulation of intracortical inhibition in human primary motor cortex. Journal of neurophysiology. 2013;110(5):1158–1166.

36. Roopun AK, Middleton SJ, Cunningham MO, et al. A beta2-frequency (20-30 Hz) oscillation in nonsynaptic networks of somatosensory cortex. Proceedings of the National Academy of Sciences of the United States of America. 2006;103(42):15646–15650.

37. Lacey MG, Gooding-Williams G, Prokic EJ, et al. Spike firing and IPSPs in layer V pyramidal neurons during beta oscillations in rat primary motor cortex (M1) in vitro. PloS one. 2014;9(1):e85109.

38. Picazio S, Veniero D, Ponzo V, et al. Prefrontal control over motor cortex cycles at beta frequency during movement inhibition. Current biology : CB. 2014;24(24):2940–2945.

39. Kopell N, Kramer MA, Malerba P, Whittington MA. Are different rhythms good for different functions? Frontiers in human neuroscience. 2010;4:187.

40. Kopell N, Ermentrout GB, Whittington MA, Traub RD. Gamma rhythms and beta rhythms have different synchronization properties. Proceedings of the National Academy of Sciences of the United States of America. 2000;97(4):1867–1872.

41. van Kerkoerle T, Self MW, Dagnino B, et al. Alpha and gamma oscillations characterize feedback and feedforward processing in monkey visual cortex. Proceedings of the National Academy of Sciences of the United States of America. 2014;111(40):14332–14341.

42. Rouiller EM, Babalian A, Kazennikov O, Moret V, Yu XH, Wiesendanger M. Transcallosal connections of the distal forelimb representations of the primary and supplementary motor cortical areas in macaque monkeys. Experimental brain research. 1994;102(2):227–243.

43. Murase N, Duque J, Mazzocchio R, Cohen LG. Influence of interhemispheric interactions on motor function in chronic stroke. Annals of neurology. 2004;55(3):400–409.

44. Stancak A, Jr., Feige B, Lucking CH, Kristeva-Feige R. Oscillatory cortical activity and movement-related potentials in proximal and distal movements. Clinical neurophysiology : official journal of the International Federation of Clinical Neurophysiology. 2000;111(4):636–650.

45. von Stein A, Sarnthein J. Different frequencies for different scales of cortical integration: from local gamma to long range alpha/theta synchronization. Int J Psychophysiol. 2000;38(3):301–313.

46. Jensen O, Tesche CD. Frontal theta activity in humans increases with memory load in a working memory task. The European journal of neuroscience. 2002;15(8):1395–1399.

47. Gevins A, Smith ME, McEvoy L, Yu D. High-resolution EEG mapping of cortical activation related to working memory: effects of task difficulty, type of processing, and practice. Cerebral cortex. 1997;7(4):374–385.

48. Rizzolatti G, Camarda R, Fogassi L, Gentilucci M, Luppino G, Matelli M. Functional organization of inferior area 6 in the macaque monkey. II. Area F5 and the control of distal movements. Experimental brain research. 1988;71(3):491–507.

49. Buffalo EA, Fries P, Landman R, Buschman TJ, Desimone R. Laminar differences in gamma and alpha coherence in the ventral stream. Proceedings of the National Academy of Sciences of the United States of America. 2011;108(27):11262–11267.

50. Barbas H. General cortical and special prefrontal connections: principles from structure to function. Annual review of neuroscience. 2015;38:269–289.

51. Derambure P, Defebvre L, Dujardin K, et al. Effect of aging on the spatio-temporal pattern of event-related desynchronization during a voluntary movement. Electroencephalography and clinical neurophysiology. 1993;89(3):197–203.

52. Hasegawa K, Kasuga S, Takasaki K, Mizuno K, Liu M, Ushiba J. Ipsilateral EEG mu rhythm reflects the excitability of uncrossed pathways projecting to shoulder muscles. Journal of neuroengineering and rehabilitation. 2017;14(1):85.

53. Horenstein C, Lowe MJ, Koenig KA, Phillips MD. Comparison of unilateral and bilateral complex finger tapping-related activation in premotor and primary motor cortex. Human brain mapping. 2009;30(4):1397–1412.

54. Berlot E, Prichard G, O’Reilly J, Ejaz N, Diedrichsen J. Ipsilateral finger representations in the sensorimotor cortex are driven by active movement processes, not passive sensory input. Journal of neurophysiology. 2019;121(2):418–426.

55. Bartos M, Vida I, Jonas P. Synaptic mechanisms of synchronized gamma oscillations in inhibitory interneuron networks. Nat Rev Neurosci. 2007;8(1):45–56.

56. Verstynen T, Ivry RB. Network dynamics mediating ipsilateral motor cortex activity during unimanual actions. J Cogn Neurosci. 2011;23(9):2468–2480.

57. Soteropoulos DS, Edgley SA, Baker SN. Lack of evidence for direct corticospinal contributions to control of the ipsilateral forelimb in monkey. The Journal of neuroscience : the official journal of the Society for Neuroscience. 2011;31(31):11208–11219.

58. Baker SN. The primate reticulospinal tract, hand function and functional recovery. The Journal of physiology. 2011;589(Pt 23):5603–5612.

59. Yao J, Dewald JPA. The Increase in Overlap of Cortical Activity Preceding Static Elbow/Shoulder Motor Tasks Is Associated With Limb Synergies in Severe Stroke. Neurorehabilitation and neural repair. 2018;32(6-7):624–634.

60. Fries P. Rhythms for Cognition: Communication through Coherence. Neuron. 2015;88(1):220–235.

