## Supplementary Material for "Coordination of multiple joints increases bilateral connectivity with ipsilateral sensorimotor cortices"

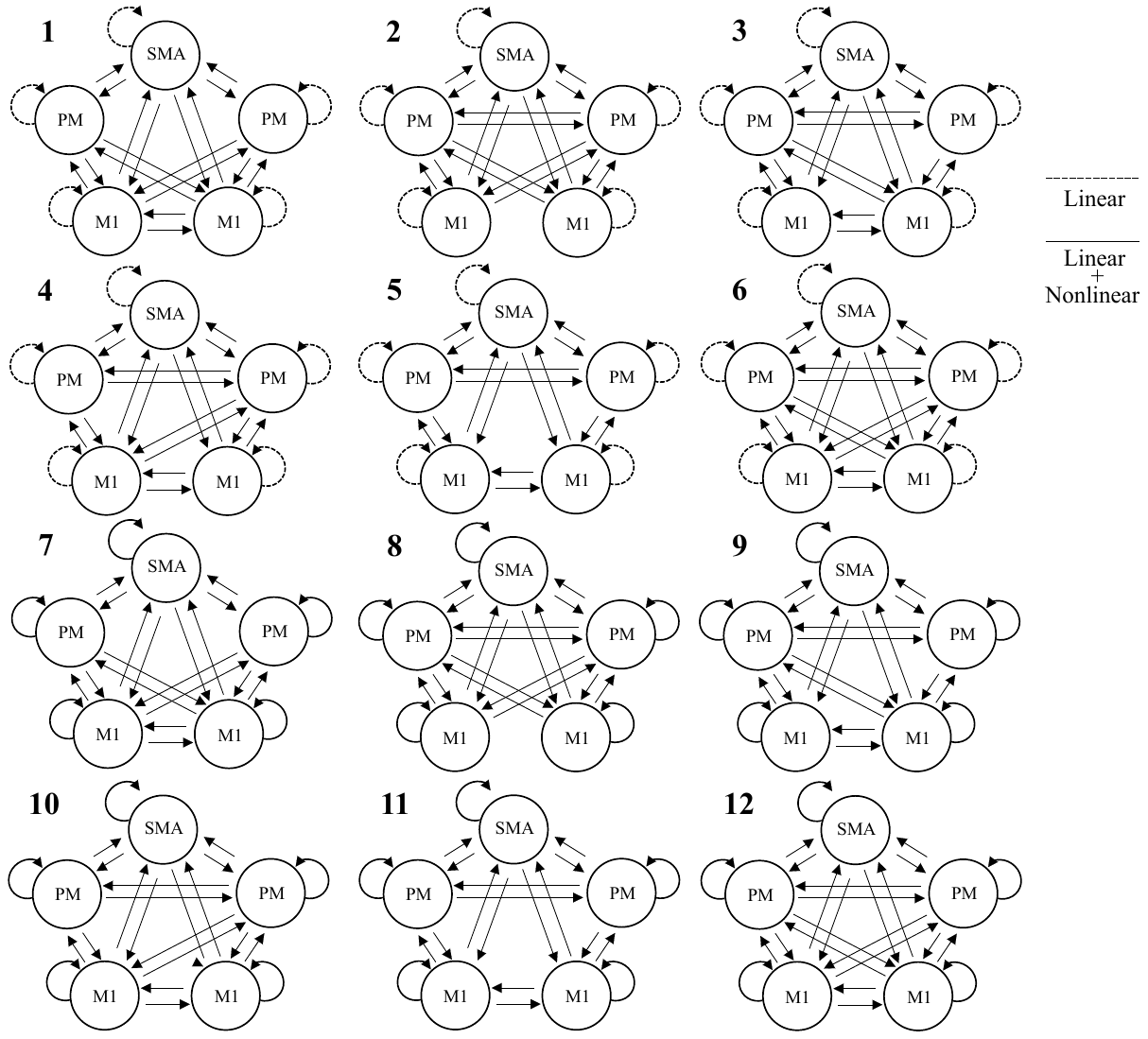

Supplementary Figure 1. Models tested for DCM analysis. Models 1-6 allow only linear intrinsic connections and both nonlinear and linear extrinsic connections, while Models 7-12 allow both nonlinear and linear intrinsic and extrinsic connections. Individual models differ in interhemispheric connections allowed between M1 and PM regions. Dashed lines indicate only linear connections allowed, while solid lines indicate both linear and nonlinear connections allowed. The lef side is the contralateral side.

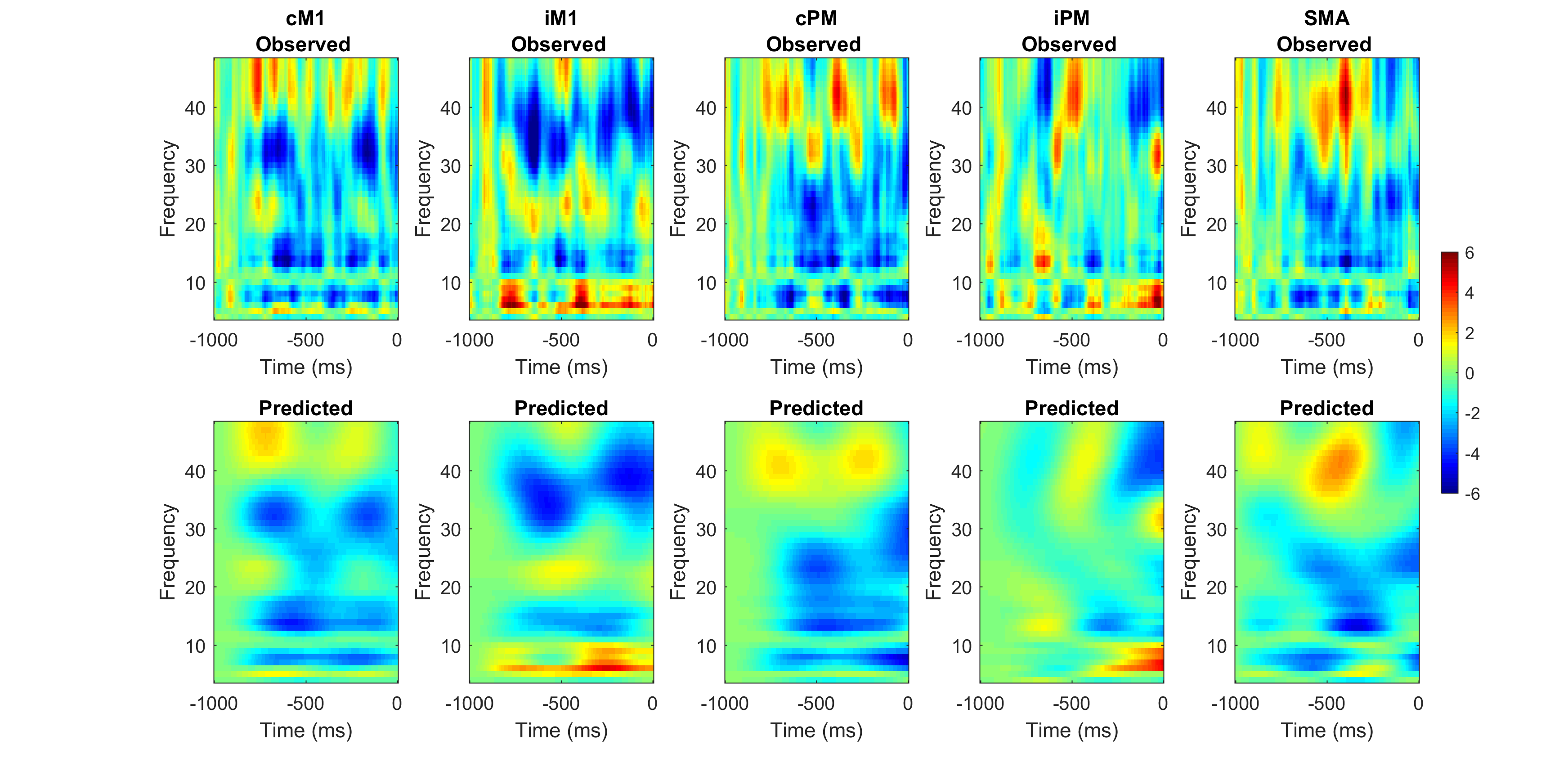

Supplementary Figure 2. The observed (top) and model-predicted (bottom) spectrograms for each region for one participant using the winning model (Model 12). Red represents an increase in power compared to baseline and blue represents a decrease in power compared to baseline. 0 ms indicates movement onset. Overall, the model explained ~80% of the original spectral variance for each condition.

Supplementary Table 1. Exceedance probabilities for each family

| Model Family | Open | Lift + Open |
| --- | --- | --- |
| Linear | 0.0001 | 0.0002 |
| Nonlinear | 0.9999 | 0.9998 |

Supplementary Table 2. Exceedance probabilities for each model

| Models | Open | Lift + Open |
| --- | --- | --- |
| 7 | 0.0001 | 0.0008 |
| 8 | 0.0001 | 0.0001 |
| 9 | 0.0003 | 0.0003 |
| 10 | 0.0000 | 0.0005 |
| 11 | 0.0001 | 0.0002 |
| 12 | 0.9994 | 0.9981 |
